# *O*-GlcNAc modifications regulate lamin A tail processing

**DOI:** 10.1101/2025.03.11.642699

**Authors:** Katherine Augspurger, Elizabeth Martin, Jason Maynard, Kevin Welle, Sina Ghaemmaghami, Al Burlingame, Barbara Panning, Abby Buchwalter

**Affiliations:** Department of Biochemistry and Biophysics, University of California San Francisco, San Francisco, United States; TETRAD Graduate Program, University of California San Francisco, San Francisco, United States; Department of Pharmaceutical Chemistry, University of University of California San Francisco, San Francisco, United States; Mass Spectrometry Resource Laboratory, University of Rochester, Rochester, New York, United States; Department of Biology, University of Rochester, Rochester, New York, United States; Cardiovascular Research Institute, University of California San Francisco, San Francisco, United States

## Abstract

Lamin A processing is highly regulated, and necessary for proper assembly of the nuclear lamina facilitating its role in nuclear structure and chromatin organization. Pre-lamin A is first farnesylated, and then a short C-terminal peptide is cleaved to produce mature lamin A. *O*-GlcNAc Transferase (OGT), a glucose sensitive post-translational modification enzyme, is a potential regulator for lamin A processing. To explore the role of OGT in lamin A biogenesis, we examined the effects of OGT levels and OGT inhibition. Variation in OGT dose or inhibition of its activity did not alter endogenous lamin A abundance or distribution. To more directly test the regulatory effects of *O*-GlcNAcylation on lamin A, we adapted a tail cleavage assay. Mutation of an OGT binding motif and *O*-GlcNAc modification sites reduced tail cleavage efficiency, suggesting that *O*-GlcNAcylation promotes lamin A processing. Our findings add to the understanding of the regulation of lamin A cleavage and identify a potential link between glucose metabolism and lamina biogenesis.

## Introduction

Lamin A is one component of the nuclear lamina, a protein meshwork that surrounds the nucleus^1,2^. The lamina connects the inner nuclear membrane to chromatin and performs two functions: providing physical rigidity to dampen external forces on the nucleus and scaffolding chromatin to influence DNA-templated processes^2^. Mutations in the nuclear lamina can lead to disease, including progeroid disorders that are characterized by rapidly advancing cardiovascular disease^3,4^.

The components of the nuclear lamina can be separated into two groups, the A-type and B-type Lamins. B-type Lamins are transcribed from their respective *LMNB1* and *LMNB2* genes, while A-type Lamins, lamin A and lamin C, are splice isoforms of the *LMNA* gene^1,4^. Uniquely, lamin A must be processed before it can be incorporated into the nuclear lamina. Mutations that perturb this processing cause Hutchinson-Gilford progeria syndrome (HGPS)^1,4^.

Lamin A processing and localization is a multi-step process. The lamin A tail is first farnesylated at a C-terminal CAAX motif, which is thought to promote association with the nuclear membrane. A C-terminal peptide is then cleaved at tyrosine 646 (Y646) by the protease ZMPSTE24^1,4^, removing the last 18 residues and the farnesylated CAAX to produce mature lamin A. A mutation that results in a loss of 50 residues around Y646 causes accumulation of farnesylated lamin A^3,4^. This incorrectly localized lamin A causes alterations in the structure of the nuclear lamina and is sufficient to cause HGPS^4,5^. This connection between aberrant lamin A biogenesis and disease highlights the importance of correct cleavage and raises the possibility that this processing can be regulated.

*O*-GlcNAc Transferase (OGT) is a potential regulator of lamin A processing^6^. OGT is an enzyme that adds a single *O*-linked sugar (*O*-GlcNAc) at Ser and Thr residues on nuclear, mitochondrial, and cytoplasmic proteins^7–9^. Lamin A is an OGT target^6^, raising the possibility that this nuclear lamina protein is regulated by this post-translational modification. Specifically, an *O*-GlcNAc site is found in close proximity to the ZMPSTE24 cut site at T643^6^. Additionally, a previously identified OGT binding motif, DNLVTRS^10^, is located immediately N-terminal to the cleavage site^6^. The proximity of the OGT binding motif and *O*-GlcNAc modification sites to the cleavage point on the lamin A tail suggests that *O*-GlcNAc modifications and/or OGT interaction may regulate lamin A processing.

We set out to test whether *O*-GlcNAcylation regulates lamin A processing using cells with natural variation in OGT levels, pharmacological inhibition of OGT, and an overexpression system to measure tail cleavage efficiency. Our results suggest that *O*-GlcNAcylation promotes lamin A processing. Exploring OGT as a novel regulator of this processing will provide insight into one molecular mechanism driving proper nuclear lamina assembly, which may be relevant to laminopathies.

## Results

### OGT and O-GlcNAc marks are more abundant in XX than in XY stem cell nuclei

*O*-GlcNAcylation of lamin A suggests OGT as a potential regulator of lamin A processing^6^. We sought to explore cell types in which there is natural variation in OGT abundance to assess how OGT may regulate lamin A. We used mouse embryonic stem cells (mESCs) because female (XX) mESCs have two active X chromosomes while male (XY) mESCs have one active X chromosome. As OGT is X-linked, XX mESCs express higher levels of OGT than males^11^. Thus, these two cell lines, that otherwise have the same developmental potential, differ in OGT abundance and provide a model system to study the effects of OGT levels on the nuclear lamina. First, we assessed OGT distribution and abundance in male and female mESCs via immunostaining and showed that OGT is concentrated in the nucleus of XX mESCs while it accumulates in the cytoplasm of XY mESCs (**Figure 1A**), as was previously shown^11^. The nuclear intensity of OGT was quantified, showing that XX mESCs have more nuclear OGT than XY mESCs (**Figure 1B**). Additionally, immunoblotting of nuclear and whole cell extracts show a greater abundance of OGT in XX mESC nuclei than in XY mESC nuclei. (**Figure 1C and 1D)**. These results establish that there is a difference in OGT abundance between XX and XY mESC nuclei, making them a useful system to study the effects of variation in OGT levels on the nuclear lamina.

**Figure 1.**
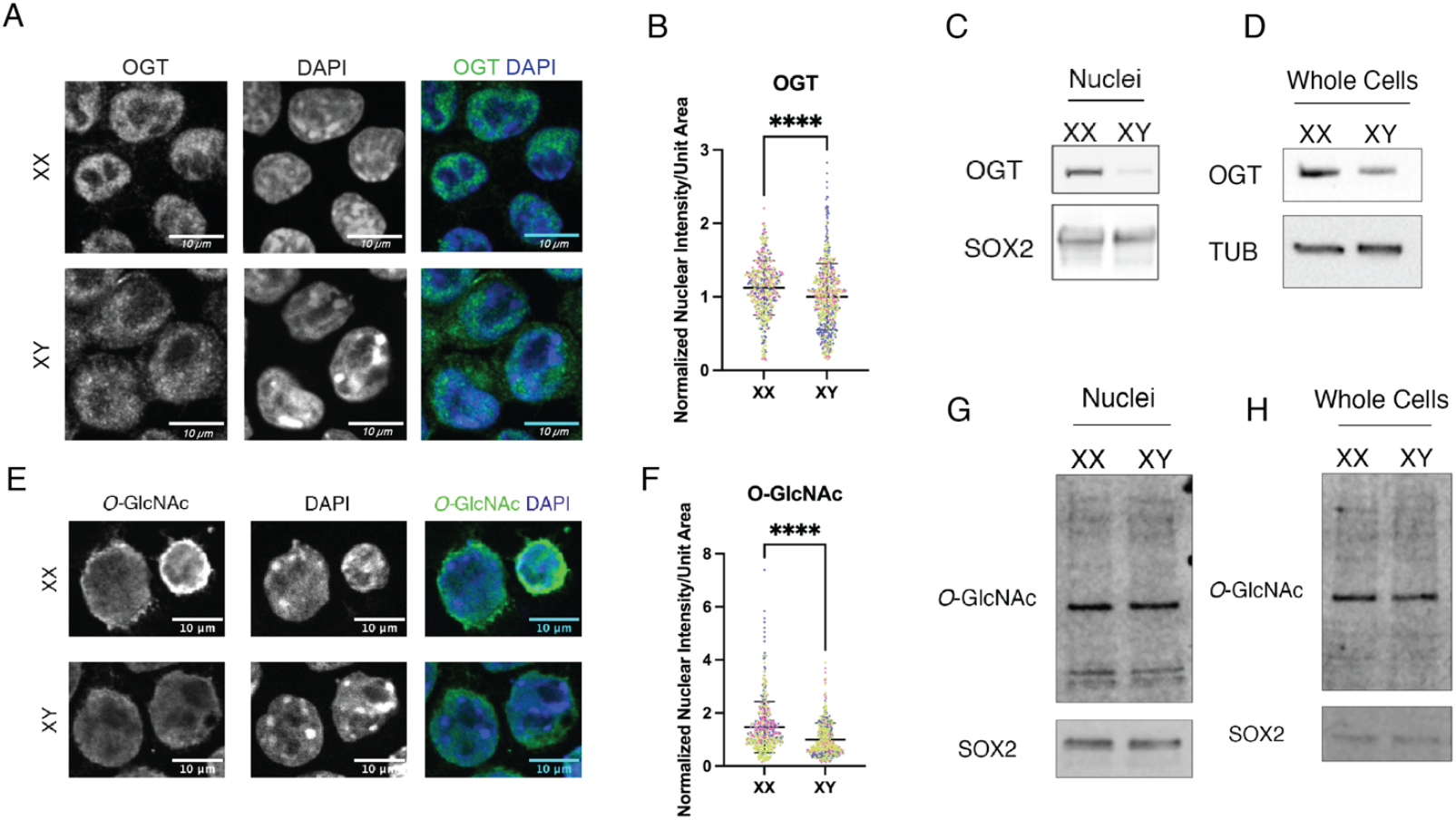
Nuclear OGT correlates with high levels of O-GlcNAcylation in XX mESCs. **A)** Confocal imaging of immunofluorescence staining of OGT on XX and XY mESCs. Representative middle slices were chosen. Scale bar is 10µm. **(B)** Quantitation of nuclear fluorescence intensity per unit area of OGT signal represented in (A) Data represents quantitation of 3 independent experiments with individual replicates indicated by distinct colors. Each experiment was normalized by the mean value of XY cells. **(C)** OGT immunoblot of XX and XY mESC nuclear extracts. 20µg nuclear protein loaded per lane. SOX2 represents loading control. **(D)** OGT immunoblot of XX and XY mESCs. 20µg whole cell lysate loaded per lane. TUB represents loading control **(E)** Confocal imaging of immunofluorescence staining of *O*-GlcNAcylation on XX and XY mESCs. Representative middle slices were chosen. Scale bar is 10µm. (**F)** Quantitation of nuclear fluorescence intensity per unit area of *O*-GlcNAc signal represented in (E). Data represents quantitation of 3 independent experiments with individual replicates indicated by distinct colors. Each experiment was normalized by the mean value of XY cells. **(G)** *O*-GlcNAc immunoblot of XX and XY mESC Nuclear extracts. 20µg nuclear extract loaded per lane. SOX2 represents loading control for stem cell nuclei. **(H)** *O*-GlcNAc immunoblot of XX and XY mESCs. 20µg whole cell protein loaded per lane. SOX2 represents loading control for stem cell nuclei. **** p < 0.0001

### O-GlcNAc enriched proteomics reveals highly O-GlcNAc modified nuclear proteins

Higher levels of OGT in the XX mESC nuclei correlate with increased nuclear *O*-GlcNAc immunostaining (**Figure 1 E and 1F**). However, a difference in total abundance of *O*-GlcNAc in nuclei and whole cells was not detectable by immunoblotting (**Figure 1G and 1H**).

To more sensitively query effects of the nuclear accumulation of OGT on protein abundance and *O*-GlcNAc modification in XX mESCs, we used unbiased multiplexed proteomics approaches.

First, we sought to correlate the number of active X chromosomes (Xs) with nuclear protein abundance. To make this comparison, we employed tandem mass tagging (TMT) coupled with LC-MS/MS using XX, XY, and XO cells (which are XX cells that have lost one X chromosome). In two separate experiments, we compared XX mESC nuclei with XY or XO mESC nuclei and calculated the Log2 fold-change of identified proteins. The Log2 fold-change values were then graphed in a correlation plot with the XX/XO value on the Y-axis and the XX/XY value on the X-axis. The correlation plots show that most proteins do not correlate with the number of X chromosomes (**Figure 2A**). The most prominent downregulated proteins are all autosomal, and some, like DNMT3a, are known to be more highly expressed in XY mESCs^12,13^. In contrast, OGT positively correlates with the number of Xs, as it is X-linked and more abundant in the XX mESC nuclei in both the XY and XO comparisons. Other highly enriched proteins in the XX cells include HMGB3, PGRC1, and EMD, all of which are X-linked and thus expected to be more abundant in the XX mESCs **(Supplementary Data 1)**. The most highly abundant autosomal protein in the XX cells compared to both XY and XO cells is lamin A/C, suggesting that lamin A/C abundance is correlated to the number of Xs.

**Figure 2.**
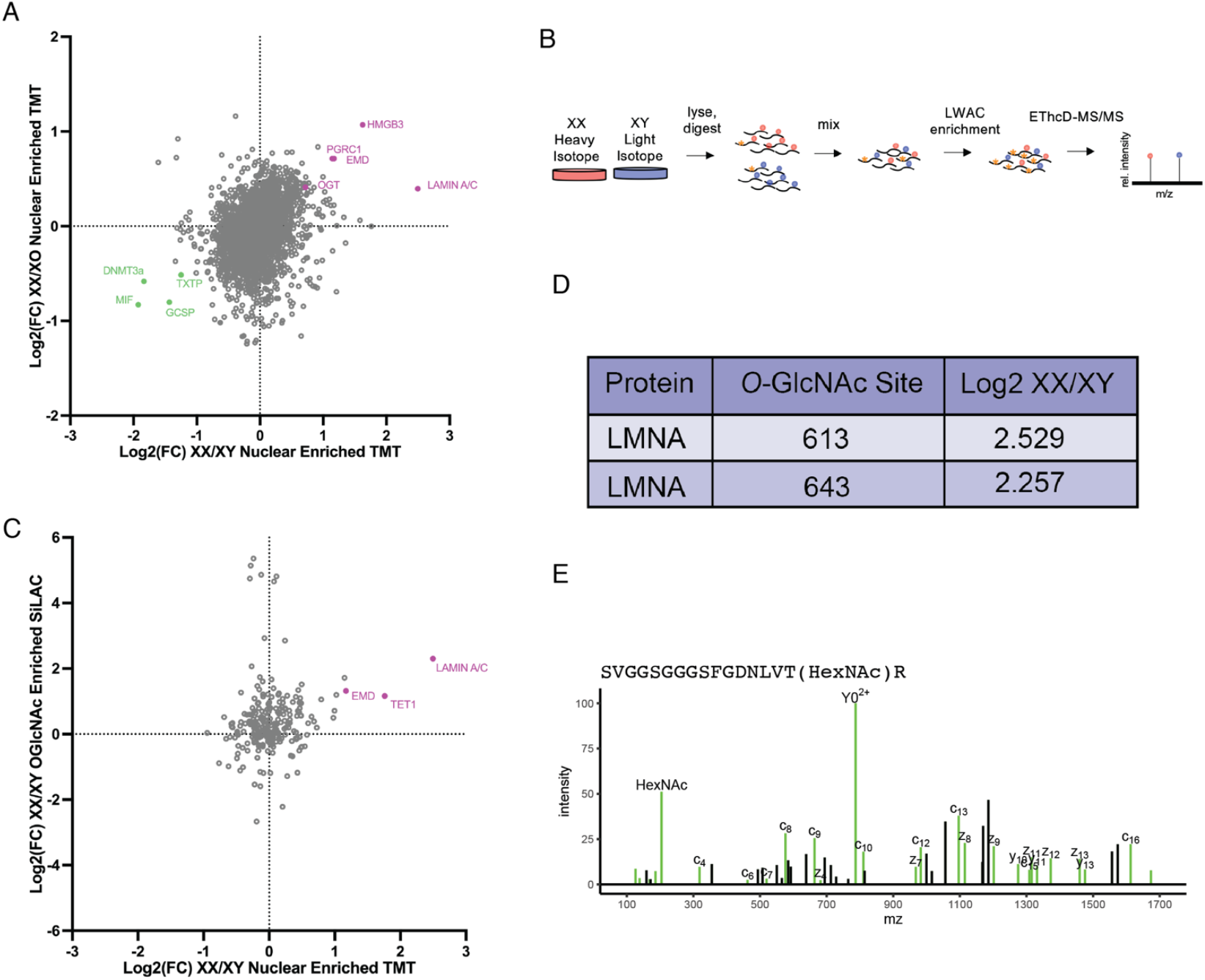
Quantitative proteomics identifies nuclear proteins that are highly O-GlcNAc modified in XX mESCs. **(A)** TMT results graphed as correlation plot of log2 fold-change of protein abundance in XX vs XY nuclei (x-axis) and XX vs XO nuclei (y-axis). **(B)** XX and XY mESCs were grown in heavy and light isotope media, respectively, and combined. Tryptic digests of combined samples were subject to lectin weak affinity chromatography (LWAC) to enrich for *O*-GlcNAcylated peptides. EThcD-MS/MS identified peptides and *O*-GlcNAc sites. **(C)** Correlation plot of log2 fold-change of protein abundance in XX vs XY nuclei (x-axis) and proteins with *O*-GlcNAc enriched peptides in XX vs XY whole cells (y-axis). **(D)** Table of *O*-GlcNAcylated peptides in lamin A and their fold change (XX/XY). **(E)** Spectrum of lamin A peptide that contains *O*-GlcNAcylation at T643.

Next, we were interested in determining which abundant proteins in the XX mESCs are also *O*-GlcNAc modified. We performed *O*-GlcNAc enriched <u>s</u>table isotope labeling using amino acids in cell culture (SILAC) mass spectrometry by growing XX and XY cell populations in heavy and light media. Labeled cells were pooled and lysed, and tryptic digests were run on a lectin weak affinity chromatography column to enrich for *O*-GlcNAcylated peptides which were quantified by EThcD-MS/MS (**Figure 2B**). In comparing the proteins enriched in XX nuclei from the TMT dataset to the identified *O*-GlcNAc modified peptides, we found X-linked OGT targets, such as emerin. In addition some autosomal proteins that have an OGT interaction motif were highly *O*-GlcNAc modified, including TET1 and lamin A (**Figure 2C, Supplementary Data 2)**. Some nuclear proteins that are highly *O*-GlcNAc modified, including nucleoporins^7^, do not exhibit increased *O*-GlcNAcylation in XX relative to XY mESC nuclei, suggesting that lamin A and TET1 are specifically X chromosome dose dependent. Analysis of individual lamin A peptides revealed two putative *O*-GlcNAc sites in the tail at S613 and T643 (**Figure 2D, 2E**). Our identification of these two sites supports previous work mapping *O*-GlcNAc marks at the same locations in liver cells^6^.

### Lamin A is highly expressed in XX mEScs

XY mESCs express low levels of lamin A^1,14^, prompting us to query the distribution of lamin A in XX mESCs. Immunofluorescence staining in XX and XY mESCs shows that lamin A is more abundant at the nuclear periphery in XX mESCs compared to the XY mESCs (**Figure 3A and B**). Immunoblot of nuclear extracts also show that both lamin A and lamin C are not detectable in XY mESC nuclei, while they are detectable in XX mESCs (**Figure 3C**). The positive correlation between the number of X-chromosomes and the amount of *O*-GlcNAcylated lamin A suggests that OGT dose may underlie lamin A abundance in XX mESCs.

**Figure 3.**
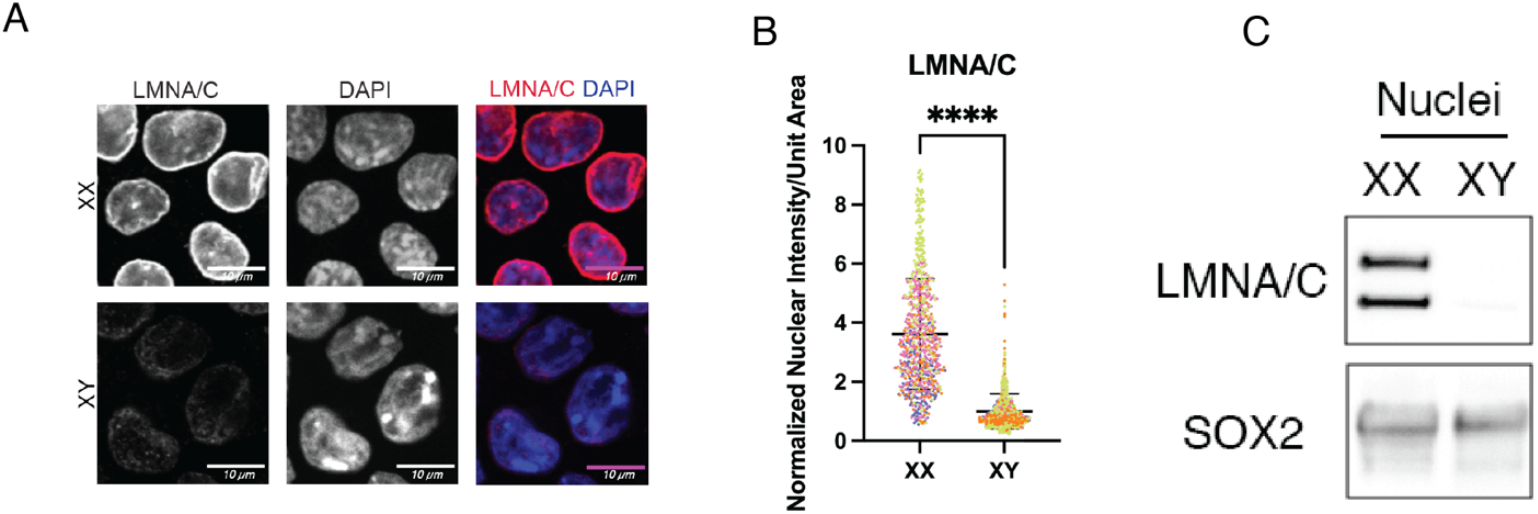
Lamin A is abundant in XX mESCs. **(A)** Confocal imaging of immunofluorescence staining of lamin A/C on XX and XY mESCs. Scale bar is 10µm. **(B)** Quantitation of nuclear fluorescence intensity per unit area of lamin A/C signal represented in (A). Data represents quantitation of 4 independent experiments with individual replicates indicated by distinct colors. Each experiment was normalized by the mean value of XY cells. **(C)** Lamin A/C immunoblot of XX and XY mESC nuclear extracts. 20µg nuclear protein loaded per lane. SOX2 represents loading control for stem cell nuclei. **** p < 0.0001

### Cellular O-GlcNAc levels do not affect lamin A protein abundance or distribution

To further understand the potential regulatory relationship between lamin A and OGT, we wanted to perturb the dosage of OGT in the XX mESCs. *OGT*, however, is an essential gene and deletion of just one copy of *OGT* is embryonic lethal in mice^15^. To circumvent the challenges of genetic manipulations, we utilized a chemical OGT inhibitor, OSMI-4^16^. OSMI-4 is a competitive inhibitor of OGT, that binds the active site. Treatment with OSMI-4 results in a successful decrease in overall *O*-GlcNAcylation in XX mESCs (**Figure 4A and D**). OGT abundance, however, increases with OSMI-4 treatment (**Figure 4A and C**). OGT homeostasis is tightly regulated, and this high OGT abundance is likely due to a feedback loop that increases OGT expression when its function or expression is perturbed^17^. The overall decrease in *O*-GlcNAcylation due to OSMI-4 treatment did not cause a significant decrease in lamin A/C abundance or distribution (**Figure 4B and E**). Lamin A/C does co-localize with lamin B1, suggesting correct localization at the nuclear lamina (**Figure S1**). Whole cell immunoblots additionally show the change in abundance of OGT and *O*-GlcNAcylation but not lamin A (**Figure 4F**). These results suggest that the overall abundance of mature lamin A is not impacted by decreased OGT activity.

**Figure 4.**
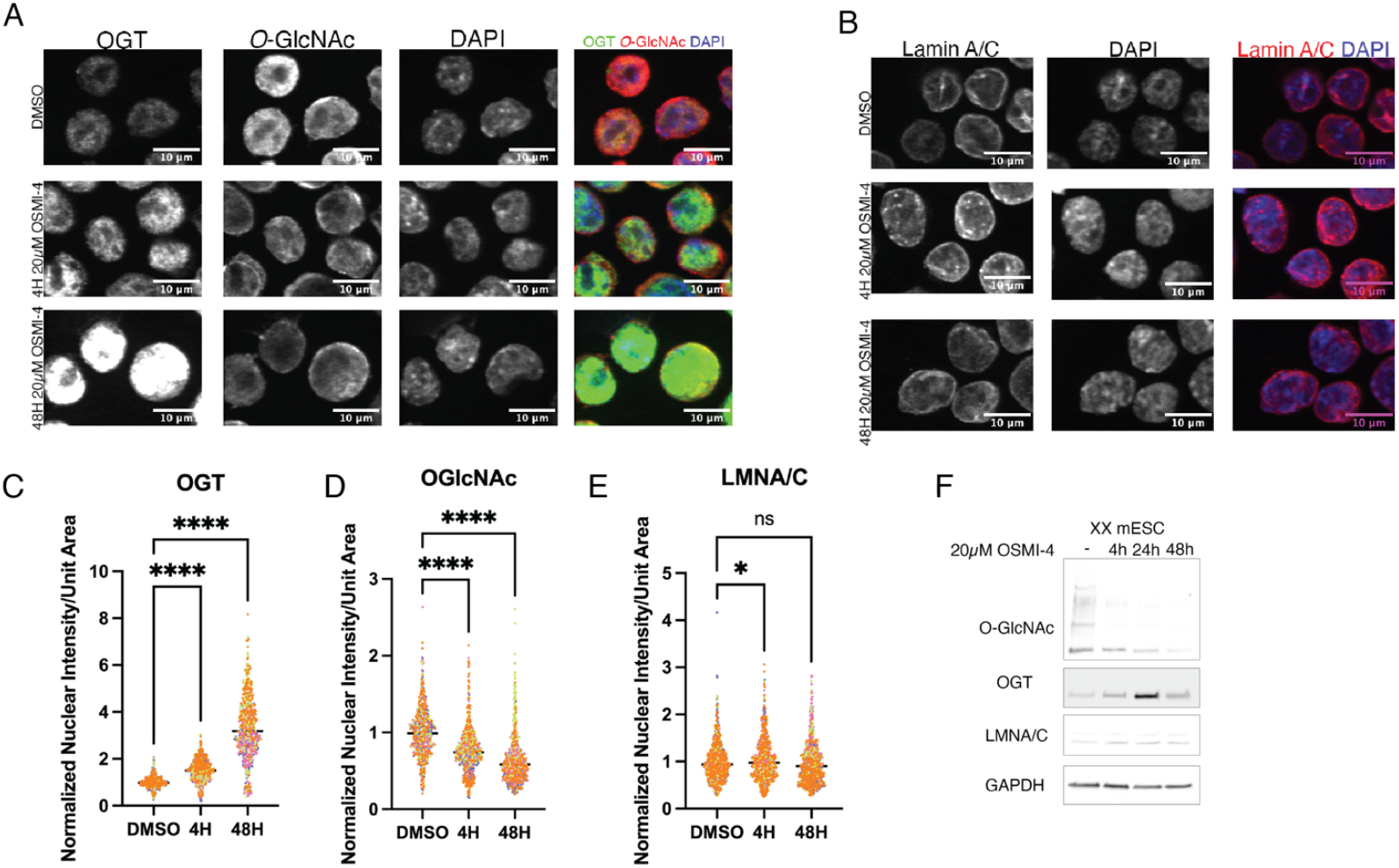
Perturbation of OGT activity in mESCs does not affect LMNA abundance or distribution. **(A)** Confocal imaging of immunofluorescence staining of OGT and O-GlcNAc on XX mESCs treated with 20µM OSMI-4 for 4 or 48 hours or with DMSO for 48 hours. Scale bar is 10µm. **(B)** Confocal imaging of immunofluorescence staining of lamin A/C on XX mESCs treated with 20µM OSMI-4 for 4 or 48 hours or with DMSO for 48 hours. Scale bar is 10µm. **(C)-(E)** Quantitation of nuclear fluorescence intensity per unit area of OGT **(C)** and *O*-GlcNAc **(D)** signal represented in (A), and lamin A **(E)** signal represented in (B). Data represents quantitation of 4 independent experiments with individual replicates indicated by distinct colors. Each experiment was normalized to the mean of the DMSO control. **(F)** Immunoblot of whole cell extracts for *O*-GlcNAc, OGT, and lamin A/C after treatment with 20µM OSMI-4 or DMSO for 4, 24, and 48 hours. GAPDH represents loading control. * p < 0.05 **** p < 0.0001 ns = not significant

### OGT promotes lamin A tail cleavage

Treatment with OSMI-4 both caused an increase in OGT abundance and a decrease in *O*-GlcNAcylation which complicates interpretation. The location of the OGT binding and *O*-GlcNAc modification sites suggests that OGT may regulate lamin A cleavage. Therefore, we directly assayed the contributions of the OGT binding motif and the *O*-GlcNAcylation sites in a lamin A tail cleavage assay ^18,19^. The T643 modification site is located just N-terminal to the ZMPSTE24 cleavage site in lamin A^1,4^. In the same region, there is also a previously identified OGT binding motif^10^ (**Figure 5A**). Doxycycline-inducible overexpression constructs encoding GFP-NLS-lamin A tails carrying point mutations that perturb *O*-GlcNAc sites and/or the OGT interaction motif (**Figure 5B**) were integrated into HEK 293T cells. Tail constructs primarily localize to the nucleus, but do not accumulate at the nuclear periphery (**Figure S2**). Roughly 24-hours after induction quantitative immunoblots were performed to resolve cleaved and uncleaved lamin A tail species (**Figure 5C**), and cleavage efficiency was quantified as the proportion of cleaved lamin A tail versus the total lamin A tail expression (cleaved + uncleaved). The cleavage efficiency was then normalized to WT, such that the WT cleavage efficiency is set at 1.0. Cleavage efficiency decreased with mutation of the Asp 639 to Ala (D639A), a mutation that disrupts the OGT binding motif^10^. Because mutation of one *O*-GlcNAcylated residue can result in modification of nearby Ser and Thr residues^20^, we mutated both T643 and S645 alone and in combination. The double mutant of T643A and S645A has a significantly lower cleavage efficiency compared to each single mutant. The triple mutant (D639A, T643A, and S645A) showed the largest decrease in cleavage efficiency with an average of 48% of the lamin A tail construct properly cleaved (**Figure 5D and E**). It was previously reported that the D639A, T643A, and S645A mutations have negligible effects on lamin A tail cleavage efficiency^21^. However, these experiments were done in a “humanized” yeast system that expresses ZMPSTE24 but does not express OGT. In total, these experiments indicate that both the OGT binding motif and the *O*-GlcNAc modified amino acids play a role in regulating the cleavage efficiency of the lamin A tail.

**Figure 5.**
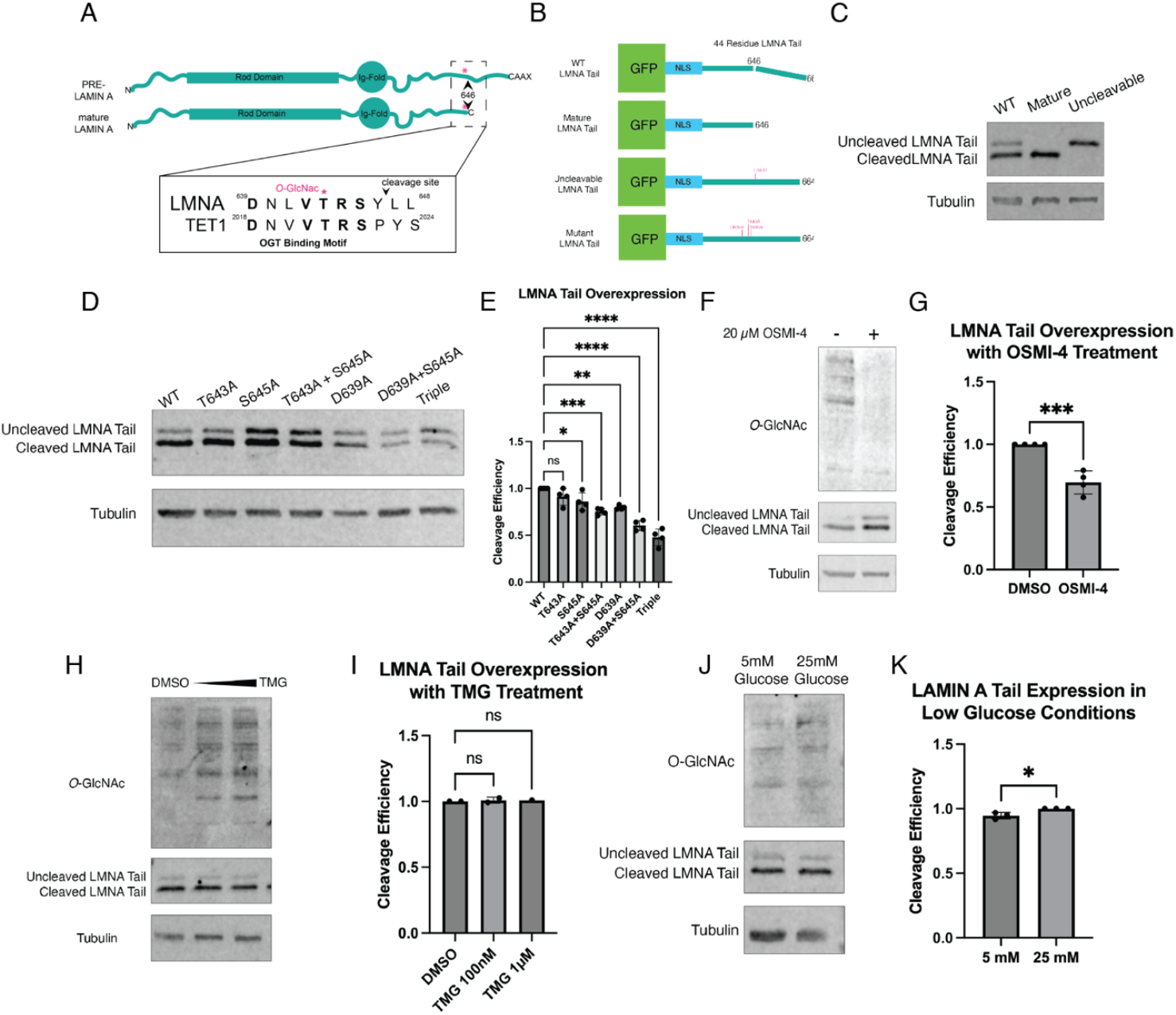
O-GlcNAc modified residues promote lamin A tail processing. (A) Diagram of lamin A tail showing the location of the T643 *O*-GlcNAc site and the OGT binding motif. (B) Diagram of lamin A tail overexpression constructs and location of point mutations in relation to the cleavage site. **(C)** GFP immunoblot of control tail cleavage constructs demonstrating size difference of cleaved and uncleaved lamin A tail. Mature lamin A tail consists of a construct that ends at Y646, mimicking the C-terminus of mature lamin A. Uncleavable mutant contains L647R mutation that fully prevents cleavage of the lamin A tail. **(D)** Quantitative GFP immunoblot of lamin A tail point mutation constructs. Tail expression induced with 1µg/mL dox for approximately 24hrs. 20µg whole cell protein loaded per lane. **(E)** Quantitation of cleavage efficiency from (D). Band intensity is normalized to tubulin loading control and background subtraction calculated based on median background intensity above and below each band. Cleavage efficiency is calculated by dividing the intensity of the cleaved LMNA tail band over the total intensity of the cleaved and uncleaved LMNA tail bands and normalized to WT. Data represents quantitation of 3 independent experiments. **(F)** Quantitative immunoblot of WT lamin A tail treated with 20µM OSMI-4 or DMSO. **(G)** Quantitation of cleavage efficiency of OSMI-4 treated lamin A tails in (F) Data represents quantitation of 3 independent experiments. **(H)** Quantitative immunoblot of WT lamin A tail treated with 100nM and 1µM TMG or DMSO **(I)** Quantitation of cleavage efficiency of TMG treated lamin A tails in (H). Data represents quantitation of 3 independent experiments. **(J)** Quantitative immunoblot of WT lamin A tail grown in media with 5mM glucose and 25 mM Glucose for 48 hours. **(K)** Quantitation of cleavage efficiency of cells in low and high glucose media in (J). Data represents quantitation of 3 independent experiments. * p < 0.05, ** p < 0.01, *** p < 0.001, **** p < 0.0001, ns = not significant

### Overall cellular O-GlcNAc levels affects lamin A cleavage efficiency

While our mutagenesis experiments revealed the importance of specific amino acids to lamin tail cleavage efficiency, these experiments cannot distinguish between the effects of the amino acid substitution and post-translational modification. To query the extent to which lamin A tail cleavage may be regulated by *O*-GlcNAc, we performed the cleavage assays with the addition of the OSMI-4 inhibitor. Treatment with OSMI-4 for 48hrs decreased both overall *O*-GlcNacylation and the cleavage efficiency of the lamin A tail constructs; OSMI-4 treated cells had a normalized cleavage efficiency of 70% (**Figure 5F and G**). We also queried the effects of increasing *O*-GlcNAcylation by treating the cells with Thiamet-G (TMG), an inhibitor of *O*-GlcNAcase (OGA), the sole enzyme that removes *O*-GlcNAcylation^7,20^. TMG treatment increased overall *O*-GlcNacylation but did not affect overall cleavage efficiency (**Figure 5H and I**), suggesting that the WT cells cleave lamin A tails at a high rate that is not influenced by additional *O*-GlcNAcylation.

Next, we wanted to test the response of the cleavage assay in a system that shows more physiological relevance. Glucose is a necessary component for synthesis of UDP-GlcNAc, OGT’s substrate, thus decreasing cellular glucose levels also decreases overall *O*-GlcNAcylation^20^ To explore the relationship between glucose levels and lamin A cleavage efficiency, WT lamin A tail construct cell lines were grown in 5 mM (low) glucose or 25 mM (high) glucose media. Cells grown in low glucose show a small but significant decrease in both overall *O*-GlcNAcylation and tail cleavage **(Figure 5J and K)**. This small decrease in cleavage efficiency may be due to the partial effect of reducing glucose on *O*-GlcNAc abundance. Together with the OSMI-4 inhibitor results, our glucose manipulation results indicate that *O*-GlcNAcylation promotes tail cleavage. These results provide evidence of a new input into control of lamin A processing.

## Discussion

Our results indicate that OGT may play a role in lamin A processing. Our data show that mutations to OGT binding and *O*-GlcNAc modification sites, OGT inhibition, and decreased glucose levels each reduce lamin A tail cleavage efficiency. However, perturbations to OGT dose do not appear to affect endogenous lamin A abundance and distribution.

In comparing lamin A abundance in XX vs XY mESCs, we have demonstrated that lamin A is more abundant in XX mESCs. In XY mESCs, lamin A expression ranges from undetectable to very low^1,14^. In contrast, we find lamin A is easily detected in XX mESCs, suggesting a correlation between X chromosome number and lamin A abundance. Lamin A is the non-X-linked protein that is most significantly enriched in XX nuclei in our quantitative proteomic data. However, in the XX vs XO dataset the relative enrichment of lamin A is lower, suggesting input from the Y chromosome may also play a role in lamin A expression. This difference in lamin A abundance highlights the importance of investigating sex difference at a cellular level as well as a whole organism level. Elucidating this relationship between sex chromosomes and lamin A may provide insight into the molecular mechanisms that control lamin A abundance.

In addition to finding more lamin A in XX mESCs, we also detected more *O*-GlcNAcylated lamin A, suggesting the overall increase in lamin A may underlie the increased abundance of the modified form. We identified two specific *O*-GlcNAc modification sites in the lamin A tail. The S613 modification is in a previously described “sweet spot”^6^ among other Ser and Thr residues. The second *O*-GlcNAc site we identified, T643, is located just N-terminal to the ZMPSTE24 cleavage site at Y646. The proximity of T643 to the cleavage site suggests that this *O*-GlcNAc site may play a regulatory role in lamin A processing.

Additional highly *O*-GlcNAc modified proteins were detected in our mass spectrometry datasets. One such protein is Emerin, which localizes to the nuclear periphery and interacts with lamin A^22–24^. Consistent with a prior report, we mapped *O*-GlcNAc sites in Emerin in the unstructured, Ser-rich lamin A binding domain^25^. As Emerin is X-linked^22^, the high abundance of *O*-GlcNAcylation detected in XX vs XY mESCs in the SILAC dataset may be the result of increased expression of Emerin in the XX mESCs.

We tested whether the high levels of OGT in XX mESC nuclei were linked to the high levels of lamin A by pharmacologically inhibiting OGT. Treatment of XX mESCs with OSMI-4 did not affect lamin A abundance or localization at the nuclear periphery. In HGPS, misprocessing of lamin A leads to nuclei with drastic morphological defects, including nuclear blebbing, aggregated nuclear pores, as well as loss of peripheral heterochromatin^5,26^.

However, we do not see these morphological changes when we inhibit OGT in the XX mESCs, suggesting that lamin A biogenesis is not detectably affected by this inhibition or that incorporation of incorrectly processed lamin A has no effect in XX mESCs. In some cell types, lamin A is a long-lived protein with slow turnover^27^. This long half-life in combination with our lack of tools to detect processing intermediates may complicate our ability to meaningfully query the effects of OGT inhibition on lamin A processing.

To more directly test lamin A processing, we examined the effects of mutation of OGT interaction and *O*-GlcNAcylation sites using an established tail cleavage assay^18,19^.

Introduction of mutations in the *O*-GlcNAc sites and/or the OGT binding motif in the lamin A tail reduced cleavage efficiency. In yeast, the same point mutants do not cause a large decrease in cleavage efficiency^21^. Yeast does not express OGT^28,29^, suggesting that the effects of mutations in human cells may in part be due to the presence of OGT. Consistent with the role of OGT, treatment with OSMI-4 also decreased the cleavage efficiency.

Additionally, growth in low glucose media which results in a decrease in abundance of the OGT co-factor UDP-GlcNAc and a concomitant decrease in *O*-GlcNAcylation^20^, caused a slight but significant decrease in processing. The decreased cleavage efficiency due to mutations, OGT inhibition, and low cellular glucose suggests that while OGT binding and *O*-GlcNAc modifications are not necessary for lamin A tail cleavage, their presence on the tail enhances cleavage.

The mechanism behind this regulation remains to be understood. It is possible that the *O*-GlcNAc modifications stabilize the interaction between lamin A and its protease ZMPSTE24. Another possibility could be that the presence of OGT and the *O*-GlcNAc marks stabilized the lamin A tail in a conformation that promotes ZMPSTE24 cleavage. A major limitation to these experiments is the use of the tagged overexpression system to assess lamin A cleavage. While this overexpression system allows us to test many point mutations, we do not retain the larger biological context of the full-length protein. Perturbing *O*-GlcNAc sites or the OGT interaction domain on the endogenous protein in OGT expressing cells would provide more insight into the mechanism linking OGT to lamin A processing.

OGT acts as a link between the proteins it modifies and metabolic regulation, as glucose is a necessary component to synthesis of OGT’s substrate UDP-GlcNAc^20^. UDP-GlcNAc levels, and thus *O*-GlcNAcylation levels, are known to fluctuate in metabolic and neurodegenerative disease^7,20^. *O*-GlcNAc modification of lamin A could act as a metabolism linked regulator of lamin A processing, suggesting a model where the integrity of the nuclear lamina may be directly affected by these metabolic disorders. Lamin A processing efficiency is also known to decrease with age^30^. However, *O*-GlcNAcylation increases with age^31,32^, perhaps in response to decreased processing efficiency. Overall, this study provides evidence for the metabolic regulation of the nuclear lamina, which may provide new insights into disease and aging.

## Data Availability

The mass spectrometry proteomics data have been deposited to the ProteomeXchange Consortium via the PRIDE^33^ partner repository with the dataset identifier PXD061673 for TMT data and identifier PXD061724 for *O*-GlcNAc enriched SILAC data.

## Acknowledgements

Data for this study were acquired on the CSU-22 Spinning Disc Confocal Microscope at the Center for Advanced Light Microscopy at UCSF. Cell lines generated for this study were sorted on a BD Aria Fusion in the Laboratory for Cell Analysis at UCSF. We’d like to thank current and past members of both the Buchwalter Lab and Panning Lab, Danica Fujimori, Jeremy Reiter, and Geeta Narlikar for valuable ideas and discussion. This work was financially supported by the Dr Miriam & Sheldon G. Adelson Medical Research Foundation (A. Burlingame), the Howard Hughes Medical Institute (A. Burlingame), NIH GM128431-02 (B. Panning), Chan Zuckerberg Biohub (A. Buchwalter), and by the UCSF Program for Breakthrough Biomedical Research (A. Burlingame, B. Panning, and A. Buchwalter).

## Methods

### Cell Culture Conditions

LF2 and E14 mESCs were routinely passaged in serum+ LIF mESC media (500 mL KO-DMEM, 10% FBS, 2 mM L-glutamine, 1X non-essential amino acids, 0.1 mM 2-mercaptoethanol and recombinant leukemia inhibitory factor) on gelatin coated tissue-culture dishes. HEK293T cells were maintained in DMEM supplemented with 10% FBS and 1X PenStrep. OSMI-4 (Sellek) or Thiamet G (Medchem Express) was added 24 hours post passage, and cells were harvested 4 hours or 48 hours later. To induce lamin A constructs, 1 µg/mL Doxycycline was added for 24 hours.

### Immunostaining

mESCs were Cytospun onto Octospot slides at 800 rpm for 3 minutes then fixed in 4% PFA for 10 minutes and washed with PBST. Fixed cells were permeablized in PBS + 0.01% TritonX-100 for 10m at room temperature and rinsed in PBST. Slides were then incubated in IF blocking buffer (10% Goat Serum, 100 µL fish skin gelatin, PBST) for 1 hour at room temperature. Slides were incubated in primary antibodies (*Supplementary Methods Table 1)* for 1 hour at room temperature and then washed 3 time in PBST. Slides incubated in fluorescent conjugated secondary antibody (*Supplementary Methods Table 2)* for 1 hour at room temperature in the dark. Slides were again washed 3 times in PBST, with the addition of DAPI in the second wash. Probed slides were mounted with prolong gold and stored at 4ºC. All primary and secondary antibodies used are listed in Methods Table 1 and Table 2.

### Image Acquisition and Analysis

Fixed and immunostained slides were imaged on a Nikon Ti inverted fluorescence microscope with CSU-22 spinning disk confocal with a 60x, 1.4 numerical aperture, oil objective. 16-bit ND2 images were acquired in 0.3 µm step Z-stacks using an EMCCD camera. Specific Z-slices were chosen and cropped for figures in FIJI. Nuclear intensity analysis was performed in CellProfiler. CellProfiler outputs were processed in r and graphed with Prism. P-values calculated with unpaired t-test in Prism.

### Nuclei Isolation

Cells were harvested, washed in PBS, and resuspended in Nuclei Preparation buffer I (320 mM sucrose, 10 mM Tris (pH 8.0), 3 mM CaCl_2_, 2 mM Mg(OAc)_2_, 0.1 mM EDTA, 0.1% Triton X-100, 1X protease inhibitors, and 1X phosphatase inhibitors) and dounce-homogenized on ice until >95% of nuclei stained by Trypan blue. Two volumes of nuclear preparation buffer II (2.0 M sucrose, Tris (pH 8.0), 5 mM Mg(OAc)_2_, 5 mM DTT, 20 μM Thiamet G, 1X protease inhibitors, and 1X phosphatase inhibitors) were added to the nuclei suspension. Nuclei were pelleted by ultracentrifugation at 130,000 × g at 4°C for 45 min. Pelleted nuclei were washed with cold PBS and stored at −80°C.

### Immunoblotting

Cells were released from tissue-culture dish and washed 1X in PBST. For endogenous lamin A blots, cell pellets were resuspended in urea lysis buffer (8 M Urea, 75 mM NaCl, 50 mM Tris pH 8.0, 1X Complete protease inhibitor tablet, and Phosphatase Inhibitor) and sonicated using a probe sonicator in 10s on/30s off cycle for 3 cycles at 30% amplitude. Total protein was quantified with Pierce BCA Assay Kit (Fisher). Urea lysates were run on a BioRad 4-20% MiniProtean-TGX gel and transferred to PVDF (0.2 µm, BioRad). Blots were then blocked in 5% milk and incubated in primary antibody (*Supplementary Methods Table 1*) at 4 ºC overnight. Blots were incubated with HRP-Conjugated secondary antibody (*Supplementary Methods Table 2*) and visualized with Pierce™ ECL Western Blotting Substrate.

For lamin A tail blots, 293T pellets were lysed in RIPA (150mM sodium chloride, 1% NP-40, 0.5% sodium deoxycholate, 0.1% sodium dodecyl sulfate, 50mM Tris, pH 8.0, 20 µM Thiamet-G, 1X protease inhibitors, 1X phosphatase inhibitors) for 20m while rotating at 4 ºC. Lysates were run on a hand-poured 12% acrylamide gel and transferred to PVDF. Blots were then blocked in 5% milk (Carnation) and incubated in primary antibody (*Supplementary Methods Table 1*) at 4 ºC overnight. Blots were incubated with fluorescently conjugated secondaries (*Supplementary Methods Table* 2) and visualized with LiCor imaging system. Band intensity was quantified in imageStuido, further analyzed with r, and graphed with Prism. P-values calculated with one-way Anova test.

### Sample Preparation for TMT

Nuclei preparations were lysed in TMT-optimized buffer (8M urea, 75 mM NaCl, 50 mM HEPES pH 8, 20µM Thiamet-G, 1X protease inhibitors, and 1X phosphatase inhibitors) and sonicated in 10s on/30s off cycle for 3 cycles at 30% amplitude. Total protein was quantified with Pierce BCA Assay Kit (Fisher). 200µg total protein was set aside for mass spectrometry. Samples were diluted to 1 mg/mL in 5% SDS, 100 mM TEAB, and 25 µg of protein from each sample was reduced with dithiothreitol to 2 mM, followed by incubation at 55°C for 60 minutes. Iodoacetamide was added to 10 mM and incubated in the dark at room temperature for 30 minutes to alkylate the proteins. Phosphoric acid was added to 1.2%, followed by six volumes of 90% methanol, 100 mM TEAB. The resulting solution was added to S-Trap micros (Protifi), and centrifuged at 4,000 x g for 1 minute. The S-Traps containing trapped protein were washed twice by centrifuging through 90% methanol, 100 mM TEAB. 1 µg of trypsin was brought up in 20 µL of 100 mM TEAB and added to the S-Trap, followed by an additional 20 µL of TEAB to ensure the sample did not dry out. The cap to the S-Trap was loosely screwed on but not tightened to ensure the solution was not pushed out of the S-Trap during digestion.

Samples were placed in a humidity chamber at 37°C overnight. The next morning, the S-Trap was centrifuged at 4,000 x g for 1 minute to collect the digested peptides. Sequential additions of 0.1% TFA in acetonitrile and 0.1% TFA in 50% acetonitrile were added to the S-trap, centrifuged, and pooled. Samples were frozen and dried down in a Speed Vac (Labconco) prior to TMTpro labeling.

### TMT Labeling

Samples were reconstituted in TEAB to 1 mg/mL, then labeled with TMTpro 16plex reagents (Thermo Fisher) following the manufacturers protocol. Briefly, TMTpro tags were removed from the -20°C freezer and allowed to come to room temperature, after which acetonitrile was added. Individual TMT tags were added to respective samples, and the reaction was allowed to occur at room temperature for 1 hour. 5% hydroxylamine was added to quench the reaction, after which the samples for each experiment were combined into a single tube. Since we performed quantitation on the unlabeled peptides, 0 day samples were added to four of the unused channels, increasing the signal for the unlabeled peptides. TMTpro tagged samples were frozen, dried down in the Speed Vac, and then desalted using homemade C18 spin columns to remove excess tag prior to high pH fractionation.

### High pH Fractionation for TMT

Homemade C18 spin columns were activated with two 50 µL washes of acetonitrile via centrifugation, followed by equilibration with two 50 µL washes of 0.1% TFA. Desalted, TMTpro tagged peptides were brought up in 50 µL of 0.1% TFA and added to the spin column. After centrifugation, the column was washed once with water, then once with 10 mM ammonium hydroxide. Fractions were eluted off the column with centrifugation by stepwise addition of 10 mM ammonium hydroxide with the following concentrations of acetonitrile: 2%, 3.5%, 5%, 6.5%, 8%, 9.5%, 11%, 12.5%, 14%, 15.5%, 17%, 18.5%, 20%, 21.5%, 27%, and 50%. The sixteen fractions were concatenated down to 8 by combining fractions 1 and 9, 2 and 10, 3 and 11, etc. Fractionated samples were frozen, dried down in the Speed Vac, and brought up in 0.1% TFA prior to mass spectrometry analysis.

### Mass Spectrometry for TMT

Peptides from each fraction were injected onto a homemade 30 cm C18 column with 1.8 um beads (Sepax), with an Easy nLC-1200 HPLC (Thermo Fisher), connected to a Fusion Lumos Tribrid mass spectrometer (Thermo Fisher). Solvent A was 0.1% formic acid in water, while solvent B was 0.1% formic acid in 80% acetonitrile. Ions were introduced to the mass spectrometer using a Nanospray Flex source operating at 2 kV. The gradient began at 3% B and held for 2 minutes, increased to 10% B over 7 minutes, increased to 38% B over 94 minutes, then ramped up to 90% B in 5 minutes and was held for 3 minutes, before returning to starting conditions in 2 minutes and re-equilibrating for 7 minutes, for a total run time of 120 minutes. The Fusion Lumos was operated in data-dependent mode, with both MS1 and MS2 scans acquired in the Orbitrap. The cycle time was set to 3 seconds. Monoisotopic Precursor Selection (MIPS) was set to Peptide. The full scan was done over a range of 400-1500 m/z, with a resolution of 120,000 at m/z of 200, an AGC target of 4e5, and a maximum injection time of 50 ms. Peptides with a charge state between 2-5 were picked for fragmentation. Precursor ions were fragmented by higher-energy collisional dissociation (HCD) using a collision energy of 38% and an isolation width of 1.0 m/z. MS2 scans were collected with a resolution of 50,000, a maximum injection time of 105 ms, and an AGC setting of 1e5. Dynamic exclusion was set to 45 seconds.

### Data Analysis for TMT

Raw data was searched using the SEQUEST search engine within the Proteome Discoverer software platform, version 2.4 (Thermo Fisher), using the Uniprot mouse database (downloaded January 2020). Trypsin was selected as the enzyme allowing up to 2 missed cleavages, with an MS1 mass tolerance of 10 ppm, and an MS2 mass tolerance of 0.025 Da. Carbamidomethyl on cysteine, and TMTpro on lysine and peptide N-terminus were set as a fixed modification, while oxidation of methionine was set as a variable modification. Percolator was used as the FDR calculator, filtering out peptides which had a q-value greater than 0.01. Reporter ions were quantified using the Reporter Ions Quantifier node, with an integration tolerance of 20 ppm, and the integration method being set to “most confident centroid”. Protein abundances were calculated by summing the signal to noise of the reporter ions from each identified peptide, while excluding any peptides with an isolation interference of greater than 40%, or SPS matches less than 65%. Further calculations were done in R and graphed in Prism.

### Sample Preparation for SILAC

XX and XY mESCs were grown under standard conditions using DMEM for SILAC, 10% dialyzed FBS, 2mM glutamine, 1X non-essential amino acids, 0.1mM b-mercaptoethanol, and recombinant leukemia inhibitory factor. XX mESCs were grown in light isotopes, L-Arginine HCl and L-Lysine-HCl. XY mESCs were grown in heavy isotopes, L-Lysine-2HCl (13C6, 15N2) and L-Arginine-HCl (13C6, 15N4) supplemented with 200 mM proline to avoid arginine-to-proline conversion. Cells were trypsinized, washed twice with cold PBS and then sonicated in 67 mM ammonium bicarbonate containing 8M guanidine HCl, 8X Phosphatase Inhibitor Cocktails II and III (Sigma-Aldrich), and 80 uM PUGNAc (Tocris Bioscience). Protein concentrations were estimated with bicinchoninic acid protein assay (ThermoFisher Scientific). 10 mg of each lysate were combined, reduced for 1 h at 56°C with 2.55 mM TCEP and subsequently alkylated using 5 mM iodoacetamide for 45 min at room temperature in the dark. Lysates were diluted to 1M guanidine HCl using 50 mM ammonium bicarbonate, pH 8.0, and digested overnight at 37°C with sequencing grade trypsin (ThermoFisher Scientific) at an enzyme to substrate ratio of 1:50 (w/w). Tryptic peptides were acidified with formic acid (Sigma-Aldrich), desalted using a 35 cc C18 Sep-Pak SPE cartridge (Waters), and dried to completeness using a SpeedVac concentrator (Thermo).

### Lectin Weak Affinity Chromatography for SILAC

Glycopeptides were enriched as described previously^34,35^. Briefly, desalted tryptic peptides were resuspended in 1000 μl LWAC buffer (100 mM Tris pH 7.5, 150 mM NaCl, 2 mM MgCl_2_, 2 mM CaCl_2_, 5% acetonitrile) and 100 μl was run over a 2.0 × 250-mm POROS-WGA column at 100 μl/min under isocratic conditions with LWAC buffer and eluted with a 100-μl injection of 40 mM GlcNAc. Glycopeptides were collected inline on a C18 column (Phenomenex). Enriched glycopeptides from 10 initial rounds of LWAC were eluted with 50% acetonitrile, 0.1% FA in a single 500-μl fraction, dried. LWAC enrichment was repeated for a total of three steps.

### Offline Fractionation for SILAC

Glycopeptides were separated on a 1.0 × 100 mm Gemini 3μ C18 column (Phenomenex). Peptides were loaded onto the column in 20 mM NH_4_OCH_3_, pH 10 and subjected to a gradient from 1 to 21% 20 mM NH_4_OCH_3_, pH10 in 50% acetonitrile over 1.1 mL, up to 62% 20 mM NH_4_OCH_3_, pH10 in 50% acetonitrile over 5.4 mL with a flow rate of 80 μL/min.

### SILAC Mass Spectrometry Analysis

Glycopeptides were analyzed on an Orbitrap Fusion Lumos (Thermo Scientific) equipped with a NanoAcquity UPLC (Waters). Peptides were fractionated on a 15 cm × 75 μM ID 3 μM C18 EASY-Spray column using a linear gradient from 2% to 30% solvent B over 65 min. Precursor ions were measured from 350 to 1800 m/z in the Orbitrap analyzer (resolution: 120,000; AGC: 4.0e5). Each precursor ion (charged 2–8+) was isolated in the quadrupole (selection window: 1.6 m/z; dynamic exclusion window: 30 s; MIPS Peptide filter enabled) and underwent EThcD fragmentation (Maximum Injection Time: 250 ms, Supplemental Activation Collision Energy: 25%) measured in the Orbitrap (resolution: 30,000; AGC; 5.04). The scan cycle was 3 s. Peak lists for EThcD were extracted using Proteome Discoverer 2.2. EThcD peak lists were filtered with MS-Filter, and only spectra containing a 204.0867 m/z peak corresponding to the HexNAc oxonium ion were used for database searching. EThcD data were searched against mouse and bovine entries in the SwissProt protein database downloaded on Sept 06, 2016, concatenated with a randomized sequence for each entry (a total of 22,811 sequences searched) using Protein Prospector (v5.21.1). Cleavage specificity was set as tryptic, allowing for two missed cleavages. Carbamidomethylation of Cys was set as a constant modification. The required mass accuracy was 10 ppm for precursor ions and 30 ppm for fragment ions. Variable modifications are listed in Supplemental Methods Table 3. Unambiguous PTMs were determined using a minimum SLIP score of six, which corresponds to a 5% local false localization rate^36^. Modified peptides were identified with a peptide false discovery rate of 1%. O-GlcNAc and O-GalNAc modifications were differentiated based on known protein subcellular localization and HexNAc oxonium ion fragments 138/144 ratio^34,37^. Comparison with TMT dataset was done in R and graphed in Prism.

### Generation of Inducible lamin A Tail Lines

Lamin A Tail constructs were created in an XLone-GFP plasmid^38^ by inserting a custom gBlock (IDT, *Methods Figure S4*) containing an NLS and the 44-amino acid lamin A tail into XLone-GFP with the NEB HiFi Assembly Kit. Next, point mutations were made using the NEB Site-Directed Mutagenesis Kit for primer design *Methods Table 5*) and assembly. Lamin A tail plasmids were co-transfected with PiggyBac Transposase^38^ with Lipofectamine 2000 (Fisher). Transfections incubated for 48 hours, and transfection media was replaced with Blasticidin selection media (DMEM, 10% FBS, 1X Pen/Strep, 6 µM Blasticidin). After selection, cells were induced with 1 µM doxycycline for 24 hours and then sorted for GFP expression (BD FACS Aria Fusion)

## Notes

### Competing Interest Statement

The authors have declared no competing interest.

