## Supplementary_Figures for "*O*-GlcNAc modifications regulate lamin A tail processing"

**Figure S1**

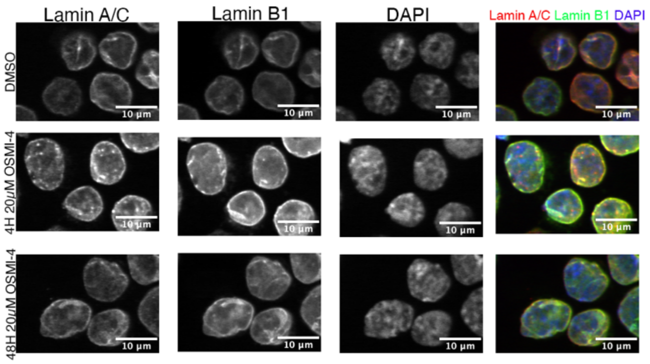


*Lamin A localization after OSMI-4 treatment*

Immunofluorescence staining of lamin A/C, lamin B1, and DAPI on XX mESCs treated with 20μM OSMI-4 for 4 or 48 hours or with DMSO for 48 hours. Scale bar is 10μm. Lamin A/C and lamin B1 co-localize at the nuclear periphery.

**Figure S2**

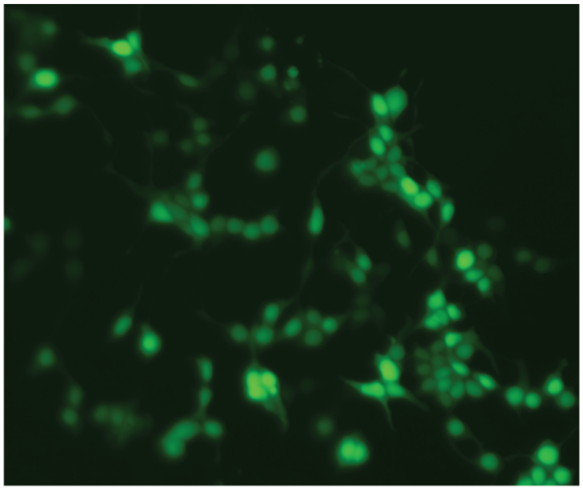

GFP-NLS-LMNA Tail (WT)

*Distribution of overexpressed lamin A tail in HEK 293T cells*

293T cells containing the WT lamin A tail construct were induced with 1μM Dox for 24hrs and GFP fluorescence was imaged on an Echo Revolve fluorescent microscope. Distribution of lamin A Tail peptide is primarily in the nucleus, with some lamin A tail also in the cytoplasm.
