## Supplementary_Methods for "*O*-GlcNAc modifications regulate lamin A tail processing"

**Supplementary Methods Table 1**

***Primary Antibodies***

| Antigen | Manufacturer | Cat. # | Dilution IF | Dilution WB |
|---|---|---|---|---|
| lamin A/C | Santa Cruz | sc-376248 | 1:100 | 1:1000 |
| *O*-GlcNAc | Abcam | ab2739 | 1:100 | 1:1000 |
| OGT | Cell Signaling Technology | D1D8Q | 1:100 | 1:1000 |
| GFP | Proteintech | pabg1 | NA | 1:1000 |
| $\alpha$-Tubulin | Sigma | T5168 | NA | 1:2000 |
| GAPDH | Genetex | GTX637966 | NA | 1:5000 |

**Supplementary Methods Table 2**

***Secondary Antibodies***

| Antibody | Manufacturer | Cat. # | Dilution |
|---|---|---|---|
| 568 AlexaFluor Goat anti Rabbit 2° | LifeTechnologies | A11011 | 1:1000 |
| 488 AlexaFluor Goat anti Mouse 2° | LifeTechnologies | A11029 | 1:1000 |
| HRP-Conjugated Goat anti Rabit 2° | VWR | RL611-1302 | 1:5000 |
| HRP-Conjugated Goat anti Mouse 2° | VWR | RL610-1302 | 1:5000 |
| Anti Rabbit 2° (DyLight680 Conjugate) | Cell Signaling Technology | 5366S | 1:5,000 – 1:20,000 |
| IRDye 800CW Goat anti Mouse 2° | LiCor | 926-32210 | 1:5,000 – 1:20,000 |

### Supplementary Methods Table 3

#### *Table of variable modifications used in the SiLAC mass spectrometry analysis*

| | |
|---|---|
| Acetyl (Protein N-term) | HexNAc4Hex6 (N) - Rare - Motif 0 N[^P][ST] |
| Acetyl+Oxidation (Protein N-term M) | HexNAc4Hex6Fuc (N) - Rare - Motif 0 N[^P][ST] |
| Gln->pyro-Glu (N-term Q) | HexNAc4Hex6Fuc2 (N) - Rare - Motif 0 N[^P][ST] |
| HexNAc (N) - Rare - Motif 0 N[^P][ST] | HexNAc4Hex6SA (N) - Rare - Motif 0 N[^P][ST] |
| HexNAc (ST) | HexNAc4Hex6SAOx2 (N) - Rare - Motif 0 N[^P][ST] |
| HexNAc2 (N) - Rare - Motif 0 N[^P][ST] | HexNAc4Hex7 (N) - Rare - Motif 0 N[^P][ST] |
| HexNAc2Fuc (N) - Rare - Motif 0 N[^P][ST] | HexNAc5Hex3 (N) - Rare - Motif 0 N[^P][ST] |
| HexNAc2Hex (N) - Rare - Motif 0 N[^P][ST] | HexNAc5Hex3Fuc (N) - Rare - Motif 0 N[^P][ST] |
| HexNAc2Hex (ST) - Rare | HexNAc5Hex3FucSA (N) - Rare - Motif 0 N[^P][ST] |
| HexNAc2Hex10 (N) - Rare - Motif 0 N[^P][ST] | HexNAc5Hex4 (N) - Rare - Motif 0 N[^P][ST] |
| HexNAc2Hex2 (N) - Rare - Motif 0 N[^P][ST] | HexNAc5Hex4Fuc (N) - Rare - Motif 0 N[^P][ST] |
| HexNAc2Hex2 (ST) - Rare | HexNAc5Hex4Fuc2 (N) - Rare - Motif 0 N[^P][ST] |
| HexNAc2Hex2Fuc (N) - Rare - Motif 0 N[^P][ST] | HexNAc5Hex4FucSA2 (N) - Rare - Motif 0 N[^P][ST] |
| HexNAc2Hex3 (N) - Rare - Motif 0 N[^P][ST] | HexNAc5Hex4NeuAc (N) - Rare - Motif 0 N[^P][ST] |
| HexNAc2Hex3Fuc (N) - Rare - Motif 0 N[^P][ST] | HexNAc5Hex4SA (N) - Rare - Motif 0 N[^P][ST] |
| HexNAc2Hex4 (N) - Rare - Motif 0 N[^P][ST] | HexNAc5Hex5 (N) - Rare - Motif 0 N[^P][ST] |
| HexNAc2Hex4Fuc (N) - Rare - Motif 0 N[^P][ST] | HexNAc5Hex5Fuc (N) - Rare - Motif 0 N[^P][ST] |
| HexNAc2Hex5 (N) - Rare - Motif 0 N[^P][ST] | HexNAc5Hex5FucSA (N) - Rare - Motif 0 N[^P][ST] |
| HexNAc2Hex5Fuc (N) - Rare - Motif 0 N[^P][ST] | HexNAc5Hex5FucSA2 (N) - Rare - Motif 0 N[^P][ST] |
| HexNAc2Hex6 (N) - Rare - Motif 0 N[^P][ST] | HexNAc5Hex5SA (N) - Rare - Motif 0 N[^P][ST] |
| HexNAc2Hex6Fuc (N) - Rare - Motif 0 N[^P][ST] | HexNAc5Hex5SA2 (N) - Rare - Motif 0 N[^P][ST] |
| HexNAc2Hex7 (N) - Rare - Motif 0 N[^P][ST] | HexNAc5Hex6 (N) - Rare - Motif 0 N[^P][ST] |
| HexNAc2Hex8 (N) - Rare - Motif 0 N[^P][ST] | HexNAc5Hex6Fuc (N) - Rare - Motif 0 N[^P][ST] |
| HexNAc2Hex9 (N) - Rare - Motif 0 N[^P][ST] | HexNAc5Hex6FucSA (N) - Rare - Motif 0 N[^P][ST] |
| HexNAc2HexFuc (N) - Rare - Motif 0 N[^P][ST] | HexNAc5Hex6FucSA2 (N) - Rare - Motif 0 N[^P][ST] |
| HexNAc3Hex3 (N) - Rare - Motif 0 N[^P][ST] | HexNAc5Hex6SA (N) - Rare - Motif 0 N[^P][ST] |
| HexNAc3Hex3Fuc (N) - Rare - Motif 0 N[^P][ST] | HexNAc5Hex6SA2 (N) - Rare - Motif 0 N[^P][ST] |
| HexNAc3Hex4 (N) - Rare - Motif 0 N[^P][ST] | HexNAc5Hex6SA3 (N) - Rare - Motif 0 N[^P][ST] |
| HexNAc3Hex4SA (N) - Rare - Motif 0 N[^P][ST] | HexNAc6Hex7FucSA (N) - Rare - Motif 0 N[^P][ST] |
| HexNAc3Hex5 (N) - Rare - Motif 0 N[^P][ST] | HexNAc6Hex7SA (N) - Rare - Motif 0 N[^P][ST] |
| HexNAc3Hex5Fuc (N) - Rare - Motif 0 N[^P][ST] | HexNAc6Hex7SA2 (N) - Rare - Motif 0 N[^P][ST] |
| HexNAc3Hex5SA (N) - Rare - Motif 0 N[^P][ST] | HexNAc7Hex6SA2 (N) - Rare - Motif 0 N[^P][ST] |
| HexNAc3Hex5SAOxSAOxAc (N) - Rare - Motif 0 N[^P][ST] | HexNAc7Hex6SA3 (N) - Rare - Motif 0 N[^P][ST] |
| HexNAc3Hex6 (N) - Rare - Motif 0 N[^P][ST] | HexNAcFuc (N) - Rare - Motif 0 N[^P][ST] |
| HexNAc3Hex6Fuc (N) - Rare - Motif 0 N[^P][ST] | HexNAcFuc (ST) - Rare |
| HexNAc3Hex6SA (N) - Rare - Motif 0 N[^P][ST] | HexNAcHex (ST) - Rare |
| HexNAc3Hex6SA2 (N) - Rare - Motif 0 N[^P][ST] | HexNAcHexFuc (ST) - Rare |
| HexNAc3Hex7 (N) - Rare - Motif 0 N[^P][ST] | HexNAcHexSA (ST) - Rare |
| HexNAc3Hex7Fuc (N) - Rare - Motif 0 N[^P][ST] | HexNAcHexSA2 (ST) - Rare |
| HexNAc4Hex3 (N) - Rare - Motif 0 N[^P][ST] | HexNAcHexSAAc (ST) - Rare |
| HexNAc4Hex3Fuc (N) - Rare - Motif 0 N[^P][ST] | HexNAcHexSAAc2 (ST) - Rare |
| HexNAc4Hex4 (N) - Rare - Motif 0 N[^P][ST] | HexNAcHexSAAcSAOxAc (ST) - Rare |
| HexNAc4Hex4Fuc (N) - Rare - Motif 0 N[^P][ST] | HexNAcHexSAOx (ST) - Rare |
| HexNAc4Hex4Fuc2 (N) - Rare - Motif 0 N[^P][ST] | HexNAcHexSAOx2 (ST) - Rare |
| HexNAc4Hex4FucSA (N) - Rare - Motif 0 N[^P][ST] | HexNAcHexSAOxAc2 (ST) - Rare |
| HexNAc4Hex4SA (N) - Rare - Motif 0 N[^P][ST] | HexNAcHexSAOxSAOxAc (ST) - Rare |
| HexNAc4Hex5 (N) - Rare - Motif 0 N[^P][ST] | HexNAcHexSASAAc (ST) - Rare |
| HexNAc4Hex5Fuc (N) - Rare - Motif 0 N[^P][ST] | HexNAcHexSASAOx (ST) - Rare |
| HexNAc4Hex5Fuc2 (N) - Rare - Motif 0 N[^P][ST] | HexNAcHexSASAOxAc (ST) - Rare |
| HexNAc4Hex5FucSA (N) - Rare - Motif 0 N[^P][ST] | HexNAcSA (ST) - Rare |
| HexNAc4Hex5FucSA2 (N) - Rare - Motif 0 N[^P][ST] | HexNAcSAOx (ST) - Rare |
| HexNAc4Hex5FucSAOx2 (N) - Rare - Motif 0 N[^P][ST] | Label:13C(6) (R) - Label 1 |
| HexNAc4Hex5SA (N) - Rare - Motif 0 N[^P][ST] | Label:13C(6)15N(2) (K) - Label 1 |
| HexNAc4Hex5SA2 (N) - Rare - Motif 0 N[^P][ST] | Met-loss (Protein N-term M) |
| HexNAc4Hex5SAOx (N) - Rare - Motif 0 N[^P][ST] | Met-loss+Acetyl (Protein N-term M) |
| HexNAc4Hex5SAOx2 (N) - Rare - Motif 0 N[^P][ST] | Oxidation (M) |
| HexNAc4Hex5SAOx3 (N) - Rare - Motif 0 N[^P][ST] | Pyro-carbamidomethyl (N-term C) |
| HexNAc4Hex5SAOxSAOxAc (N) - Rare - Motif 0 N[^P][ST] | |

### Supplemental Methods Figure 4

*<u>gBlock (GFP-NLS-lamin A Tail) for HiFi Assembly of XLone-GFP-lamin A Tail:</u>*

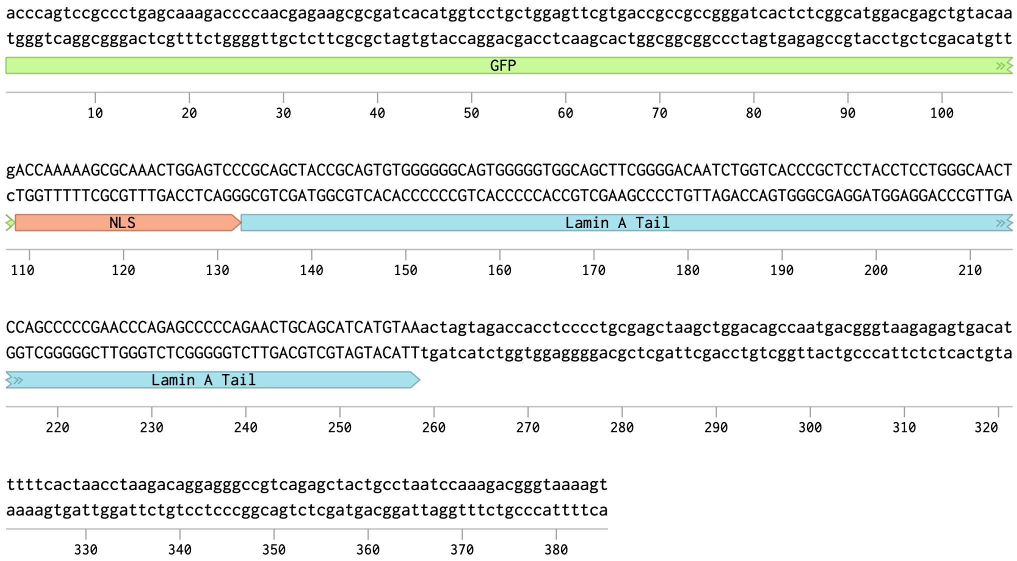

**Supplementary Methods Table 5**

Primers for lamin A tail construct and site directed mutagenesis

| Cell Line | Forward | Reverse |
|---|---|---|
| WT lamin A Tail HiFi Assembly | CGAGCTGTACAAGACCAAA<br>AAGCGCAAACTGGAGTCCC<br>GCAGCTACCGCAGTGTGG | TGCGGTAGCTGCGGGA<br>CTCCAGTTTGCGCTTTTTG<br>GTCTTGTACAGCTCGTCCATGCCGA |
| Mature lamin A Tail | TAAACTAGTAGACCACCTCCC | GTAGGAGCGGGTGACCAG |
| L647R lamin A Tail | CCGCTCCTACCGACTGGGCAACT | GTGACCAGATTGTCCCCG |
| T643A lamin A Tail | CAATCTGGTCGCCCGCTCCTACC | TCCCCGAAGCTGCCACCC |
| S645A lamin A Tail | GGTCACCCGCGCCTACCTCCTGG | AGATTGTCCCCGAAGCTGCC |
| T643A+S645A lamin A Tail | CAATCTGGTCGCCCGCGCCTACC | TCCCCGAAGCTGCCACCC |
| D639A lamin A Tail | CAGCTTCGGGGCCAATCTGGTCACCC | CCACCCCCACTGCCCCCC |
| D639A+S645A lamin A Tail | GGTCACCCGCGCCTACCTCCTGG | AGATTGGCCCCGAAGCTGC |
| Triple Mutant lamin A Tail | CAATCTGGTCGCCCGCGCCTACC | GCCCCGAAGCTGCCACCC |
